# Automated detection of Hainan gibbon calls for passive acoustic monitoring

**DOI:** 10.1101/2020.09.07.285502

**Authors:** Emmanuel Dufourq, Ian Durbach, James P. Hansford, Amanda Hoepfner, Heidi Ma, Jessica V. Bryant, Christina S. Stender, Wenyong Li, Zhiwei Liu, Qing Chen, Zhaoli Zhou, Samuel T. Turvey

## Abstract

1. Extracting species calls from passive acoustic recordings is a common preliminary step to ecological analysis. For many species, particularly those occupying noisy, acoustically variable habitats, the call extraction process continues to be largely manual, a time-consuming and increasingly unsustainable process. Deep neural networks have been shown to offer excellent performance across a range of acoustic classification applications, but are relatively underused in ecology.
2. We describe the steps involved in developing an automated classifier for a passive acoustic monitoring project, using the identification of calls of the Hainan gibbon *(Nomascus hainanus)*, one of the world’s rarest mammal species, as a case study. This includes preprocessing - selecting a temporal resolution, windowing and annotation; data augmentation; processing - choosing and fitting appropriate neural network models; and postprocessing - linking model predictions to replace, or more likely facilitate, manual labelling.
3. Our best model converted acoustic recordings into spectrogram images on the mel frequency scale, using these to train a convolutional neural network. Model predictions were highly accurate, with per-second false positive and false negative rates of 1.5% and 22.3%. Nearly all false negatives were at the fringes of calls, adjacent to segments where the call was correctly identified, so that very few calls were missed altogether. A postprocessing step identifying intervals of repeated calling reduced an eight-hour recording to, on average, 22 minutes for manual processing, and did not miss any calling bouts over 72 hours of test recordings. Gibbon calling bouts were detected regularly in multi-month recordings from all selected survey points within Bawangling National Nature Reserve, Hainan.
4. We demonstrate that passive acoustic monitoring incorporating an automated classifier represents an effective tool for remote detection of one of the world’s rarest and most threatened species. Our study highlights the viability of using neural networks to automate or greatly assist the manual labelling of data collected by passive acoustic monitoring projects. We emphasise that model development and implementation be informed and guided by ecological objectives, and increase accessibility of these tools with a series of notebooks that allow users to build and deploy their own acoustic classifiers.

## 2 Introduction

Deep learning holds enormous promise for automating the labelling of bioacoustic data. The number of applications is growing (Christin, Hervet, & Lecomte, 2019), but the majority of datasets are still labelled manually (Fairbrass et al., 2019; Kiskin et al., 2020; Pamula, Pocha, & Klaczynski, 2019), even as the rate of data collection makes this approach increasingly unsustainable. The mismatch between the potential of deep learning approaches and their actual uptake among practitioners occurs because getting models to perform as well as an experienced human is difficult. Human-like performance usually requires substantial amounts of training data or relatively stable background environments, conditions that are often absent in ecological applications. Model tuning and data manipulation is often required, and while guidelines are emerging (Patterson & Gibson, 2017; Stowell, Wood, Pamula, Stylianou, & Glotin, 2019), these can, with some justification, appear subjective and case specific. A lack of computing resources and user-friendly software can also be a barrier to entry. Case studies reporting successful applications play an important role in developing and disseminating best practices, and in discriminating between those tasks that current deep learning methods are able to automate and those they cannot. Previous applications have used convolutional neural networks (CNNs; LeCun, Bengio, and Hinton (2015)) to identify various bird (Grill & Schlüter, 2017; Kahl et al., 2017; Stowell, Wood, et al., 2019) and whale species (Bergler et al., 2019; Bermant, Bronstein, Wood, Gero, & Gruber, 2019; Jiang et al., 2019; Shiu et al., 2020), bees (Kulyukin, Mukherjee, & Amlathe, 2018; Nolasco et al., 2019), as well as anomalous acoustic events in soundscapes (Sethi et al., 2020). These have shown, for example, that a generally good approach is to represent data as spectrograms and treat the problem as an image classification one, as well as providing specialised approaches for data augmentation on spectrogram inputs, such as pitch and time shifting and introducing background noise (Bergler et al., 2019; Sprengel, Jaggi, Kilcher, & Hofmann, 2016).

Despite this, no studies report the process of applying deep learning within the scope of a typical acoustic monitoring project designed to answer a well-defined research question. Most applications are either smaller – using data collected for the purpose of testing a deep learning approach, and often written for a machine learning rather than ecological audience (e.g. Kiskin et al., 2020; Kulyukin et al., 2018); or larger – aggregating datasets across several independent studies to investigate if models generalise (Bergler et al., 2019; Shiu et al., 2020; Stowell, Wood, et al., 2019) – than most monitoring projects. In this paper we address this gap, describing the development of a classifier for identifying Hainan gibbon (*Nomascus hainanus*) calls in passive acoustic recordings collected as part of a long-term monitoring project, with the aim of providing practitioners with a realistic and relatable idea of the process, and modelling choices, involved, as well as guidelines for these choices.

The Hainan gibbon is the world’s rarest primate and one of the world’s rarest mammals, with only a single population of about 30 individuals surviving in Bawangling National Nature Reserve (BNNR), Hainan, China (Chan, Fellowes, Geissmann, & Zhang, 2005; Liu, Ma, Cheyne, & Turvey, 2020; S. Turvey et al., 2015). Improved monitoring of this population using novel methods, to understand factors affecting successful dispersal, breeding group formation and colonization of new habitat, has been identified as an urgent short-term conservation goal for the species (S. Turvey et al., 2015; Zhang et al., 2020). Gibbons call regularly to advertise territory and maintain group cohesiveness against rivals, using a complex structure consisting of short individual vocal syllables or “notes” of ca. 0.2–2.75s assembled together into longer “phrases” consisting of one to six notes, which are themselves organised into “songs” of several minutes (Deng, Zhou, & Yang, 2014). Gibbon population surveys are usually conducted by detecting this daily song using a fixed-point count survey method, whereby researchers listen opportunistically for calls at elevated listening posts (Brockelman & Srikosamatara, 1993; Kidney et al., 2016). However, this traditional monitoring approach is labour-intensive and is only conducted for discrete survey periods. Gibbons are therefore prime candidates for passive acoustic monitoring and recent studies have used data collected in this way to model occupancy (Vu & Tran, 2019) and to discriminate between individuals using spectral features (Clink, Crofoot, & Marshall, 2019; Zhou et al., 2019). All of these studies, however, have relied on an initial manual extraction of calls.

In order to develop a continuous monitoring protocol for Hainan gibbons we conducted longterm passive acoustic monitoring and developed an automated classifier able to identify whether gibbons were calling in the vicinity of a particular recorder, with the aim of establishing whether the area proximal to the recorder was occupied that day. It was therefore important to be able to detect individual gibbon calling bouts, but not necessarily to be able to discriminate every phrase made during the bout. We address issues that are important to the overall usefulness of a classifier, including deciding how much data to manually label, data augmentation, operationally meaningful definitions of classifier success, and the development of user-friendly software. Our study provides an effective new monitoring method for the world’s rarest primate, and also has wider applicability for applying deep learning to develop passive acoustic monitoring frameworks for other conservation-priority loud-call species such as cetaceans, elephants, or other primates (Crunchant, Borchers, Kuhl, & Piel, 2020).

## 3 Materials and Methods

### 3.1 Data collection

Eight Song Meter SM3 recorders (Wildlife Acoustics, Maynard, Massachusetts) were used to collect acoustic data from 1 March to 20 August 2016 within BNNR. Recorders were attached to trees at a height of approximately 1.5m in tropical evergreen forest. Four recorders were situated within the known home ranges of the four Hainan gibbon social groups existing during the study period (Groups A-D; see Bryant, Zeng, Hong, Chatterjee, and Turvey (2017)), three were situated at locations intermediate between known home ranges, and a further recorder was placed in an area where a solitary male gibbon was thought to occur (Bryant et al., 2016). They were placed at locations that were used as regular listening posts for monitoring gibbons by reserve staff (Figure 1). The peak Hainan gibbon calling period is 06:00–07:00, with calling continuing at decreasing regularity for several hours (Chan et al., 2005). Recorders were therefore set to record for eight hours each day from the time of sunrise, which varied between approximately 05:00 and 06:00 during the study period. Memory cards and batteries were changed every 40 days. Devices did not record continuously throughout the entire survey period due to logistical and technical issues; in total, survey days per recorder varied between 79 and 129 days, and roughly 6,000 hours of recordings were collected. The majority of recordings were made with a sampling rate of 9,600Hz and bit depth of 16, with isolated recordings at 28,800Hz.

**Figure 1:**
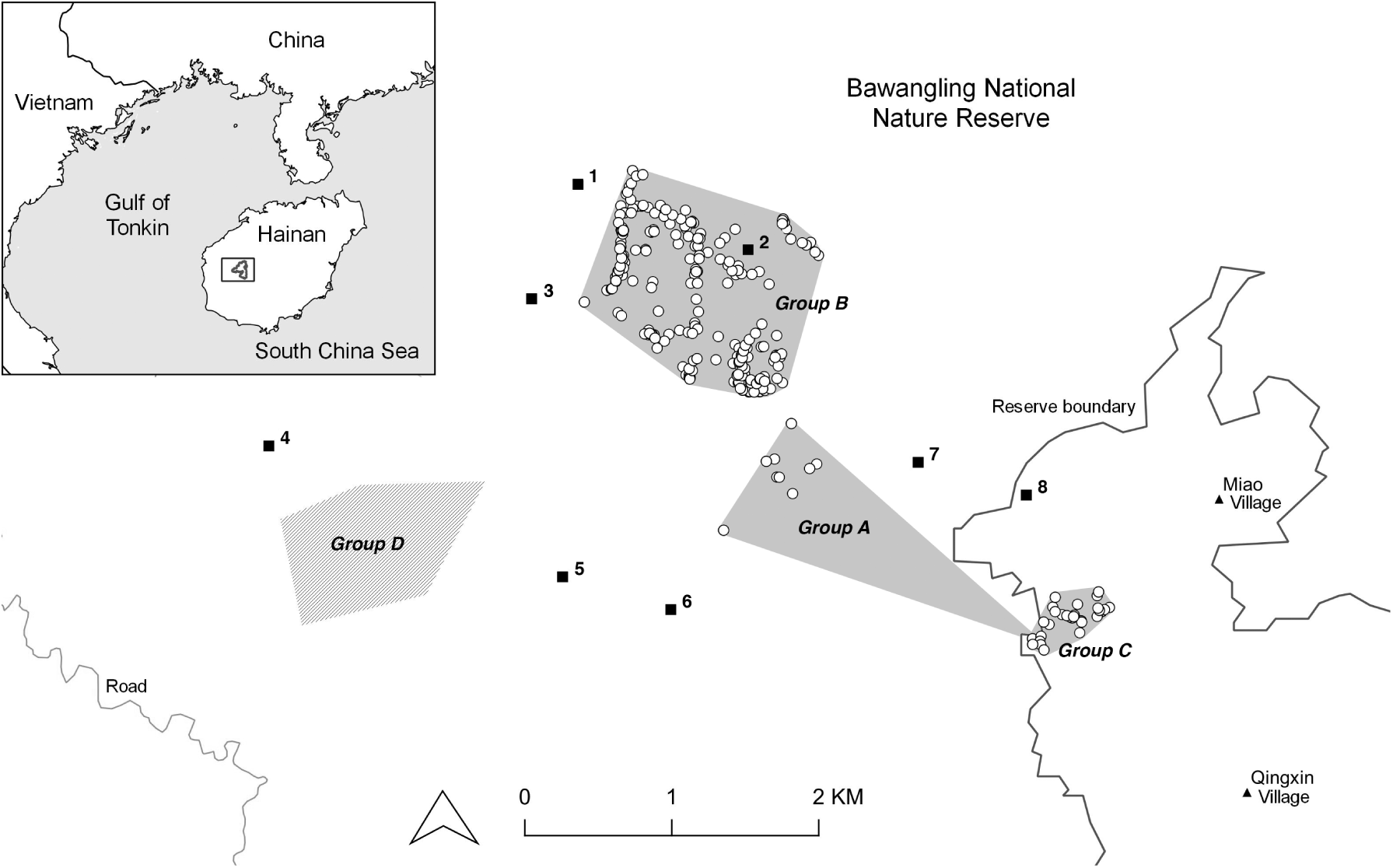
Locations of eight Song Meter SM3 recorders (labelled 1-8) used to detect gibbons in 2016 within Bawangling National Nature Reserve, Hainan, China, in relation to approximate distributions of four Hainan gibbon social groups (A-D). Mapped distributions of groups A-C are based on field data collected in 2010-2011 (see Bryant et al. (2017)); the groups all changed their location slightly between 2011 and 2016, but data on exact group locations in 2016 are unavailable. Approximate location of Group D indicated with hatching based on Bryant et al. (2016).

### 3.2 Data analysis

We manually labelled 32 eight-hour recordings by inspecting spectrograms and listening to audio using Sonic Visualiser (Cannam, Landone, & Sandler, 2010), and recording the start and end times, and the number of notes, of each observed gibbon phrase. This process yielded 1,246 gibbon phrases.

To construct the fixed-length inputs required by CNNs, we divided each eight-hour recording into segments with window length 10s and hop length 1s (starting times of consecutive 10s segments differ by 1s, Figure 2). This window length was chosen so that even the longest phrase (8s, Supplementary Material A) fits within a single segment; using a slightly longer segment length allows for potentially longer unseen phrases, and results in more positive segments after windowing. All audio was converted into mono, as done in various applications (e.g. Bergler et al., 2019; Qazi, Tabassam Nawaz, Rashid, & Habib, 2018; Stowell, Petrusková, Šálek, & Linhart, 2019). By crossreferencing the time intervals of each segment with the logged start and end times of known gibbon phrases, each segment was labelled as (a) a “presence”, if its time interval completely contained the interval of at least one labelled phrase, (b) an “absence”, if its time interval contained no part of any phrase, or (c) a “partial presence”, if its time interval intersected but did not completely contain the interval of at least one labelled phrase (Figure 2). Partial presences were excluded from further analysis.

**Figure 2:**
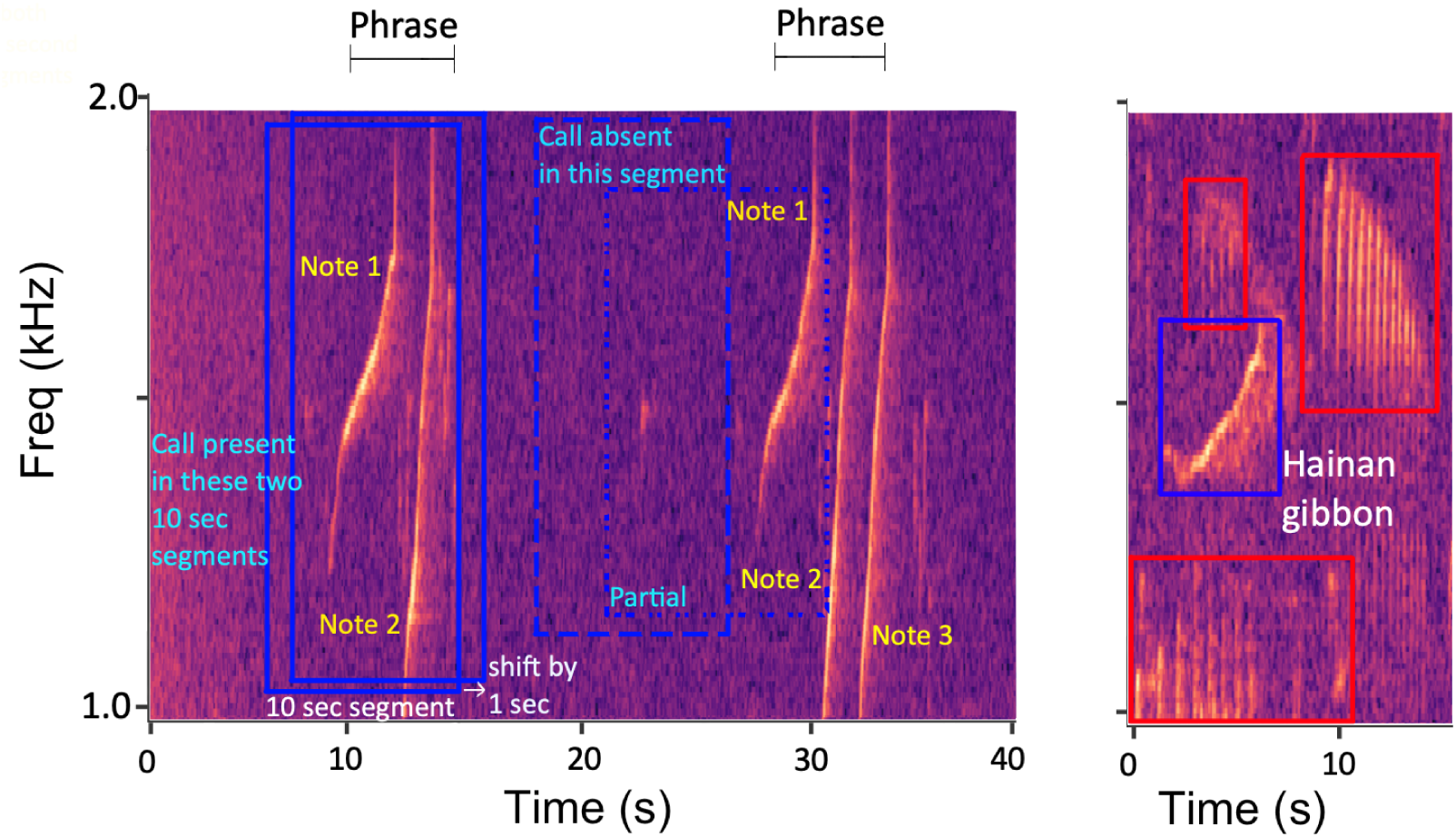
Hainan gibbon calls consist of a sequence of “phrases”, each phrase consisting of variable (typically, 1-6) “notes” and often with relatively large intervals between phrases. **Left:** a two-note phrase followed by a three-note phrase. A single calling bout may last anywhere from a few to dozens of minutes. Our model divides the recording interval into sliding 10s windows or “segments” (blue boxes), with 80% overlap between adjacent segments. Segments are classified as contained at least one full gibbon phrase (Present; solid line), a partial phrase (Partial; dotted line), or no part of a phrase (Absent; dashed line). Partial presences were excluded from further analysis, creating a two-class audio classification problem. **Right:** a gibbon phrase partially obscured by noisy background conditions, in this case other species calling (red boxes).

Preprocessed amplitudes in each 10s segment were downsampled to 4800Hz, and the downsampled inputs – each segment a time series of 48000 observations – used as inputs to the 1-D CNNs described in the next section. In addition, we converted each audio segment into a mel-scale spectrogram (Bergler et al., 2019; Huang, Acero, & Hon, 2001), to be used as an input image to a 2-D CNN, using a window size of 1,024/9,600s, a hop size of 256/9,600s, and 128 mel frequency bins with centres uniformly spaced between 1 and 2kHz, a conservative interval following Deng et al. (2014) and our own exploratory analyses. These values for chosen on the basis of preliminary investigations, although results are not particularly sensitive to these choices. The spectrogram images had a size of 128 *×* 188 pixels; larger image sizes can capture greater detail but typically require more network parameters and computation time to do so.

After processing, our dataset consisted of 5,285 segments containing at least one complete phrase. While the vast majority of segments do not contain any gibbon calls, we restricted the number of absence segments to the same number as presences, to avoid a large class imbalance. Absence segments were initially collected by randomly sampling, but we found that better results were obtained by specifically including absence segments that contained typical ambient noise, such as bird calls, rain events, and other background noises that could potentially confuse the classifier (Stowell, Petrusková, et al., 2019). Extracting these required additional manual processing of the audio data.

### 3.3 Data augmentation

Data augmentation – boosting sample sizes by adding new samples artificially created by manipulating existing ones, for example using geometric operations like translations and rotation – is commonly used to improve classifier performance, particularly when the training dataset is relatively small (Hestness et al., 2017; Sun, Shrivastava, Singh, & Gupta, 2017). We used data augmentation to create up to ten new copies of each 10s segment in both presence and absence classes. For each presence segment **x**^(*pre*)^, we randomly selected ten absence segments, 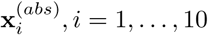. We randomly shifted the starting time of each absence segment forward by 0 *< t*_*i*_ *<* 9 seconds, with the absence segment wrapping back on itself so that it remained 10s long (Figure 3c), to obtain the shifted segment 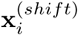. Presence segments were not shifted, as this already occurred during the windowing process used to create the original segments. Segments contain amplitude values and thus allow for arithmetic operations to be performed on them. We blended the presence segment with each shifted segment to create augmented presence segments 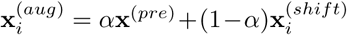, where *α* is a mixing parameter, here chosen to be 0.9 (Figure 3d). We created augmented absence segments using the same approach, i.e. combining pairs of absence segments to create a mixture of background scenes.

**Figure 3:**
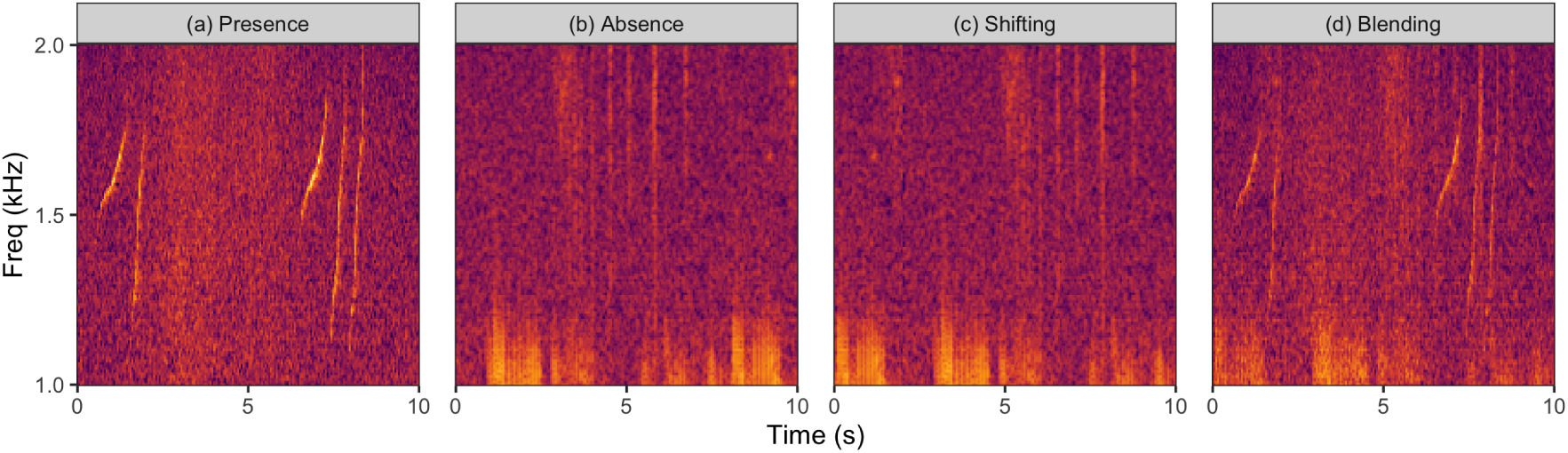
Data augmentation steps involve (a) selecting a presence segment containing a Hainan gibbon phrase, (b) randomly selecting a segment containing only background noise, (c) shifting the starting time of the absence segment forward by a random amount, here two seconds, and (d) blending together the presence and shifted absence segments.

After augmenting the original segments, we obtained 18,992 segments (9,496 presence, 9,496 absence) from 19 recordings to train the neural networks. We randomly selected 60% of the data for training (5,697 presence, 5,697 absence) and used the remaining 40% for validation (3,799 presence, 3,799 absence). Non-augmented segments from nine separate recordings (2,231 presence, 23,689 absence) were kept aside for testing.

### 3.4 Neural networks

We considered two kinds of CNN architectures: a 1-D CNN using preprocessed amplitudes of 10s segments as inputs, and a 2-D CNN that had inputs consisting of spectrogram images constructed from the preprocessed amplitudes. As we had relatively little training data by deep learning standards, we chose these networks as they use simple architectures requiring relatively few parameters. Both 1-D and 2-D CNNs use up to three convolutional layers, each followed by a max pooling layer that reduces the size of the intermediate input passed to the next layer of the network. We used 16 *×* 1 and 16 *×* 16 convolutional kernels for 1-D and 2-D CNNs, respectively. The stack of convolutional layers was followed by one or two dense layers (Figure 4). The resulting model outputs a predicted probability that the input segment (1-D or 2-D) contains at least one complete gibbon phrase.

**Figure 4:**
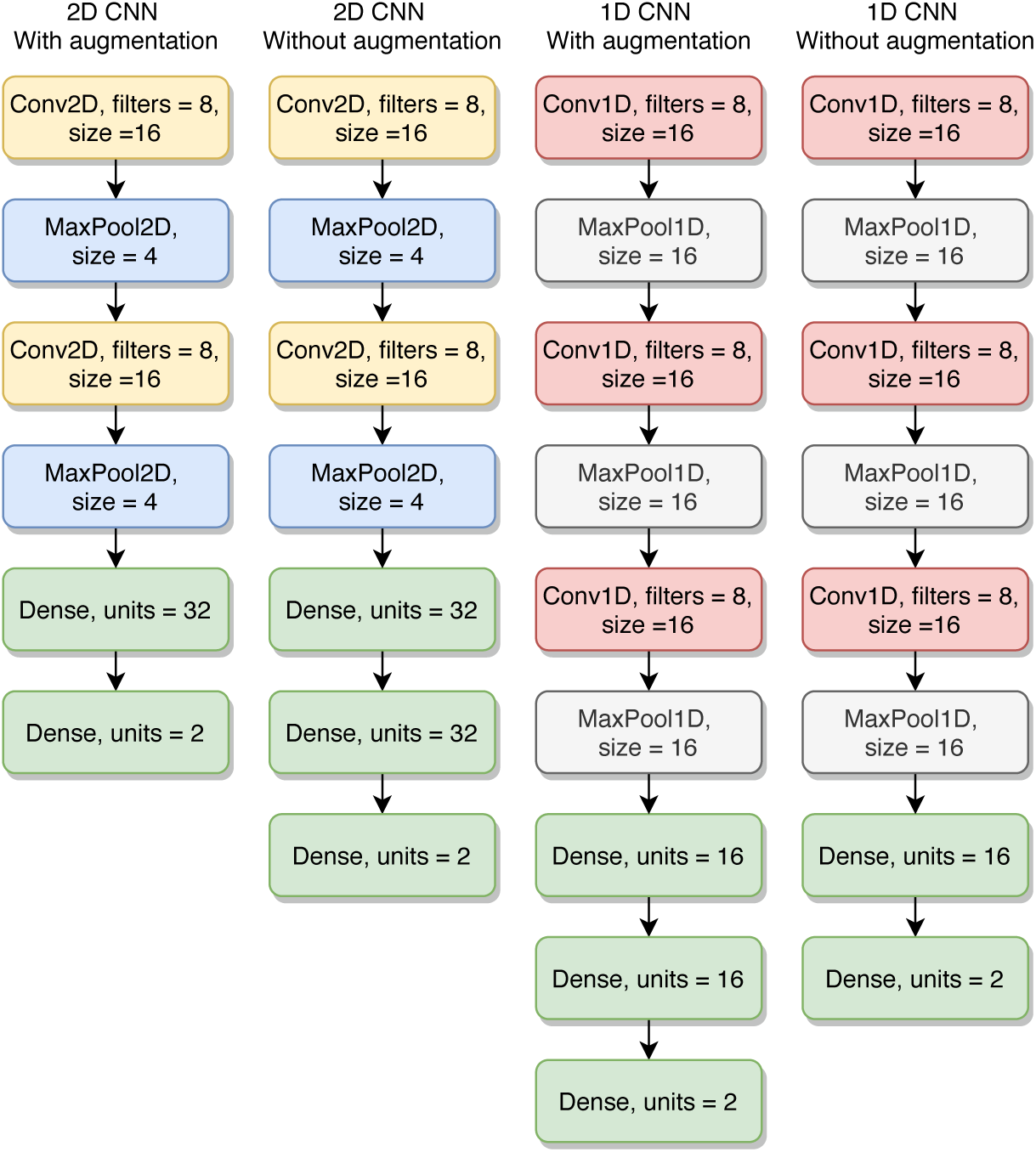
Best architectures for 1-D and 2-D CNNs, for both augmented and non-augmented training datasets. Selected architectures were those with intermediate numbers of free parameters, particularly for 2-D CNNs.

We chose model hyperparameters using a grid search over the number of convolutional (1, 2, 3) and dense (1, 2, 3) layers, nodes in each of the dense layers (8, 16, 32), filters in each convolutional layer (8, 16, 32), kernel size in each convolutional and max pooling layer (4, 8, 16), and dropout rate (0, 0.2, 0.4, 0.6). Each model was trained for 50 epochs using the Adam optimizer (Kingma & Ba, 2014) a batch size of 8 segments, and a learning rate of 0.001. Models were evaluated based on test set accuracy (proportion of all predictions that were correct), sensitivity (proportion of true positives divided by positive examples), and specificity (proportion of true negatives divided by negative examples). Optimal thresholds for converting predicted probabilities into binary classifications were those that minimized the ratio of sensitivity and false discovery rate in the validation dataset.

Models were implemented in Python 3 using the TensorFlow (Abadi et al., 2015) library with Keras (Chollet et al., 2015) for the neural network component, and the Librosa library for audio processing and spectrogram construction (McFee et al., 2020). Model training and testing was done on a machine running Ubuntu 16.04 LTS with an Intel i7-6700K CPU, 16GB of RAM, and an Nvidia GTX 1070 8GB Graphics Processing Unit. Code and analysis scripts are available online at https://github.com/emmanueldufourq/GibbonClassifier.

### 3.5 Post-processing

For an audio recording of arbitrary duration, our approach was to break that recording into overlapping 10s segments, and to use a trained CNN to output, for each segment starting at second *s* = 0, 1, 2, …, a predicted probability indicating the likelihood that at least one complete gibbon phrase is contained in the next ten seconds. These probabilities are based only on the acoustic content of their associated segments, and can give rise to biologically unrealistic call patterns. We used a post-processing step to remove isolated predicted presence segments which are highly likely to be false positives rather than actual calls, and to obtain start and end times for each predicted calling bout, to facilitate manual verification and support the main research objective of detecting and monitoring gibbon activity.

To do this, we formed connected components of presence segments that occur close together in time and in sufficient numbers that, given known gibbon call characteristics (i.e. song duration, inter-phrase duration), they are likely to be part of a single calling bout (Supplementary Material A). With presence segments arranged in temporal order, presence segment *i* is included in the same component as segment *i−*1 if they are separated by less than 200s; otherwise segment *i* begins a new component. This process allocates each presence segment to exactly one component. Components were then reviewed, and any components consisting of fewer than 20 segments (equivalent to roughly four phrases of length 5s) were removed, as were any components where the average time between consecutive presence segments in the component was greater than 10s (suggesting a “chain” of isolated presence predictions, since calls usually persist over multiple consecutive segments).

The first and last presence segment in each remaining component give the start and end times of each predicted gibbon calling bout. To evaluate the post-processing step, we mimic its intended application by assuming that all predicted bouts are passed to an observer for manual processing, and that all presence segments within the bout are subsequently identified. This approach means that post-processing accuracy measures are conditional on the use of additional, error-free manual verification.

## 4 Results

Hainan gibbon calls could be detected with a high degree of accuracy. Without post-processing, nearly 80% of segments containing gibbon calls were correctly identified, with very few false positives (Table 1). Even with false negative rates of 20% very few gibbon phrases were missed altogether, because phrases occur across multiple overlapping segments and nearly all segments incorrectly identified as absences occurred at the beginning and end of a phrase, abutted by several segments where the phrase was correctly detected (Figure 5). After post-processing, fewer than 2% of all presence segments occurred outside of predicted call bouts (Table 1), and all 20 call bouts across nine test set recordings were detected, with two predicted call bouts being false positives (Supplementary Material B). In the training set, 34 of 35 call bouts were correctly recognised with 2 false positive call bouts.

**Table 1:** Average classification accuracy and parameter settings for the best 2-D and 1-D CNN models across 72 hours of test recordings (2,231 segments containing gibbon phrases, 23,689 without). Gibbon calls can be identified with very high accuracy, and performance is improved by data augmentation and a postprocessing heuristic.

| CNN<br>+ Augmentation<br>+ Postprocessing | 2-D<br>Yes<br>Yes | 2-D<br>Yes<br>No | 2-D<br>No<br>No | 1-D<br>Yes<br>Yes | 1-D<br>Yes<br>No | 1-D<br>No<br>No |
| --- | --- | --- | --- | --- | --- | --- |
| Accuracy (Test) | 99.37% | 97.60% | 92.32% | 94.30% | 94.76% | 94.76% |
| Sensitivity (Test) | 98.30% | 77.68% | 79.65% | 54.21% | 40.98% | 25.56% |
| Specificity (Test) | 99.42% | 98.51% | 92.92% | 95.96% | 96.91% | 97.60% |
| Accuracy (Train) | 98.68% | 97.20% | 93.65% | 95.14% | 94.16% | 93.44% |
| Sensitivity (Train) | 94.84% | 80.64% | 77.85% | 69.62% | 53.42% | 24.53% |
| Specificity (Train) | 99.12% | 98.59% | 94.94% | 97.66% | 97.92% | 99.24% |
| Model Parameters | 23,922 | 23,922 | 24,978 | 2,650 | 2,650 | 2,378 |
| Train Duration (sec) | 644 | 643 | 265 | 628 | 627 | 117 |

**Figure 5:**
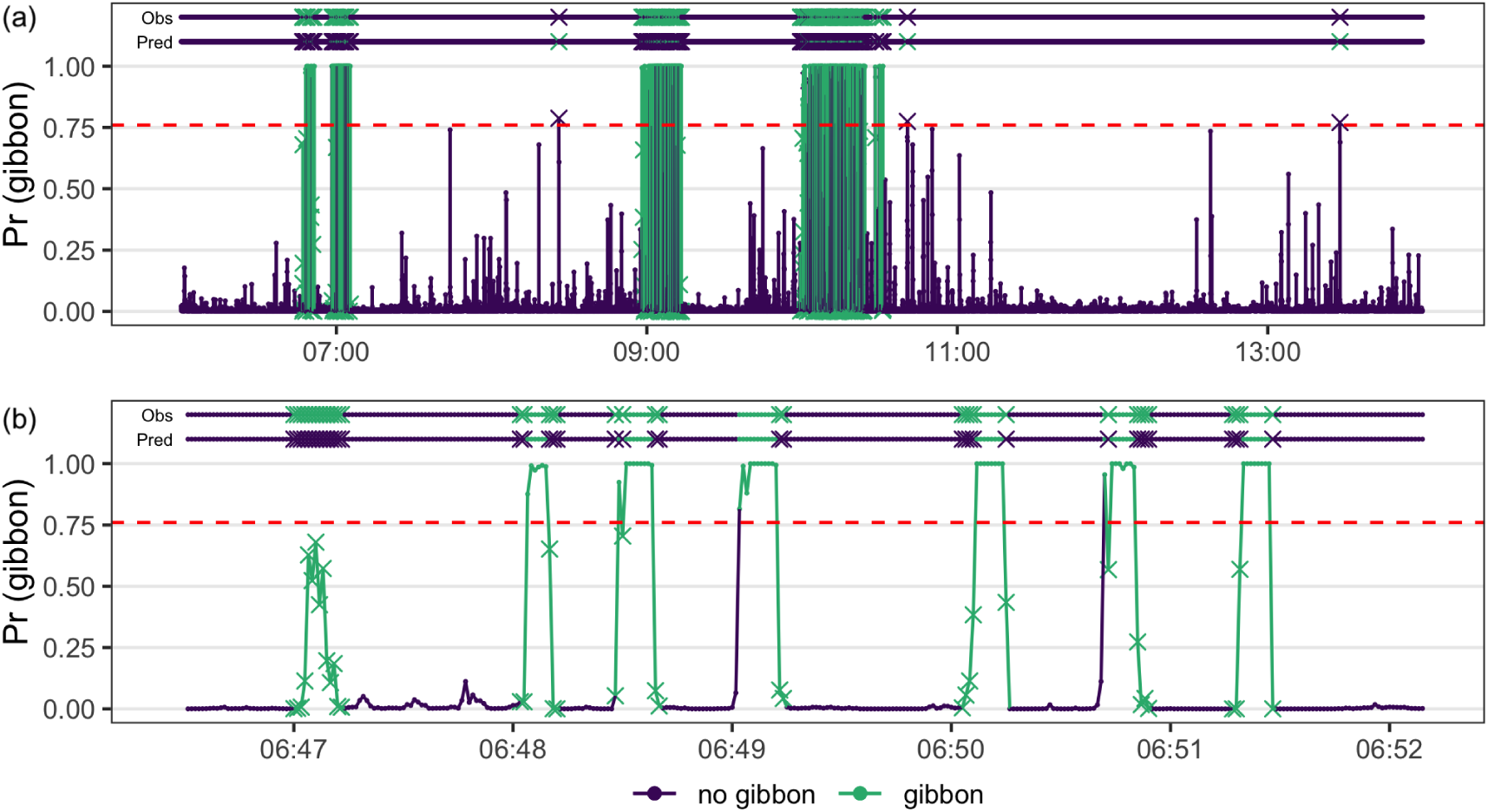
Per-second predicted probabilities that a gibbon phrase is contained within the next 10s of audio, over (a) an eight-hour file, (b) a five-minute window. Segments with predicted probabilities above an optimized threshold of 0.76 (red line) are classified as containing a gibbon phrase, with misclassifications denoted by crosses. Observed and predicted classes are plotted above the probabilities, using the same notation. Colour is used to denote the observed class. Most incorrect false negative classifications are at the beginning and end of phrases, separated by segments that correctly identify the call. In this way, nearly all phrases are clearly identified, and a practitioner can be pointed to those regions that contain calls.

The best performing approach was a 2-D CNN with both data augmentation and post-processing. Data augmentation improved specificity by 5.6%, a relative reduction in false positives of 79% but without associated relative reduction in sensitivity; post-processing further improved both sensitivity (20.6%) and specificity (0.9%, Table 1). Accuracy was substantially higher when treating the task as an image (spectrogram) classification problem than if the preprocessed acoustic data were directly used as input to a 1-D CNN. An 8 hour test file took on average 6 minutes to process of which 3 minutes 10 seconds were used for reading in the audio file and 2 minutes 42 seconds to convert to spectrograms; the remaining time was used to compute the CNN predictions.

Across the entire monitoring project, gibbon calls were detected on 71% of recording days across all locations. Gibbons were detected regularly at all locations, with recorders situated within known group or solitary home ranges detecting calls on 33–86% of recording days, and those situated between home ranges detecting calls on 46–89% of recording days. Mean durations of calling bouts per recorder varied between 24.2 and 40.8 minutes (overall mean = 29.7 minutes), with mean starting times of 06:16-07:56 am and mean finishing times of 09:12-10:15 am (Figure 6; Table 2). Calls were detected less frequently during the wet season (March-April) than the dry season (May-August), with inter-season differences varying substantially between locations (Supplementary Table C).

**Table 2:** Calling behaviour across 8 survey locations for the 161 day survey period March–August 2016. Recorders were situated within the known home ranges of the four Hainan gibbon social groups existing during the study period, at locations intermediate between known home ranges, and in an area where a solitary male gibbon was thought to occur. Locations of home ranges are indicated by numbers 1, 2, 3 and 4. 6 = solitary.

| Location | Survey days | % days calls detected | Mean calling time per day (min) | Mean start time of first bout | Mean end time of last bout |
| --- | --- | --- | --- | --- | --- |
| 1 | 87 | 70 | 24.2 | 07:34 | 09:41 |
| 2 | 90 | 46 | 29.9 | 06:58 | 09:12 |
| 3 | 103 | 82 | 31.3 | 07:30 | 10:15 |
| 4 | 105 | 86 | 26.5 | 07:44 | 09:52 |
| 5 | 79 | 33 | 29.9 | 07:31 | 09:23 |
| 6 | 103 | 79 | 24.4 | 07:56 | 10:15 |
| 7 | 129 | 89 | 30.9 | 06:53 | 09:54 |
| 8 | 105 | 65 | 40.8 | 06:16 | 10:01 |

**Figure 6:**
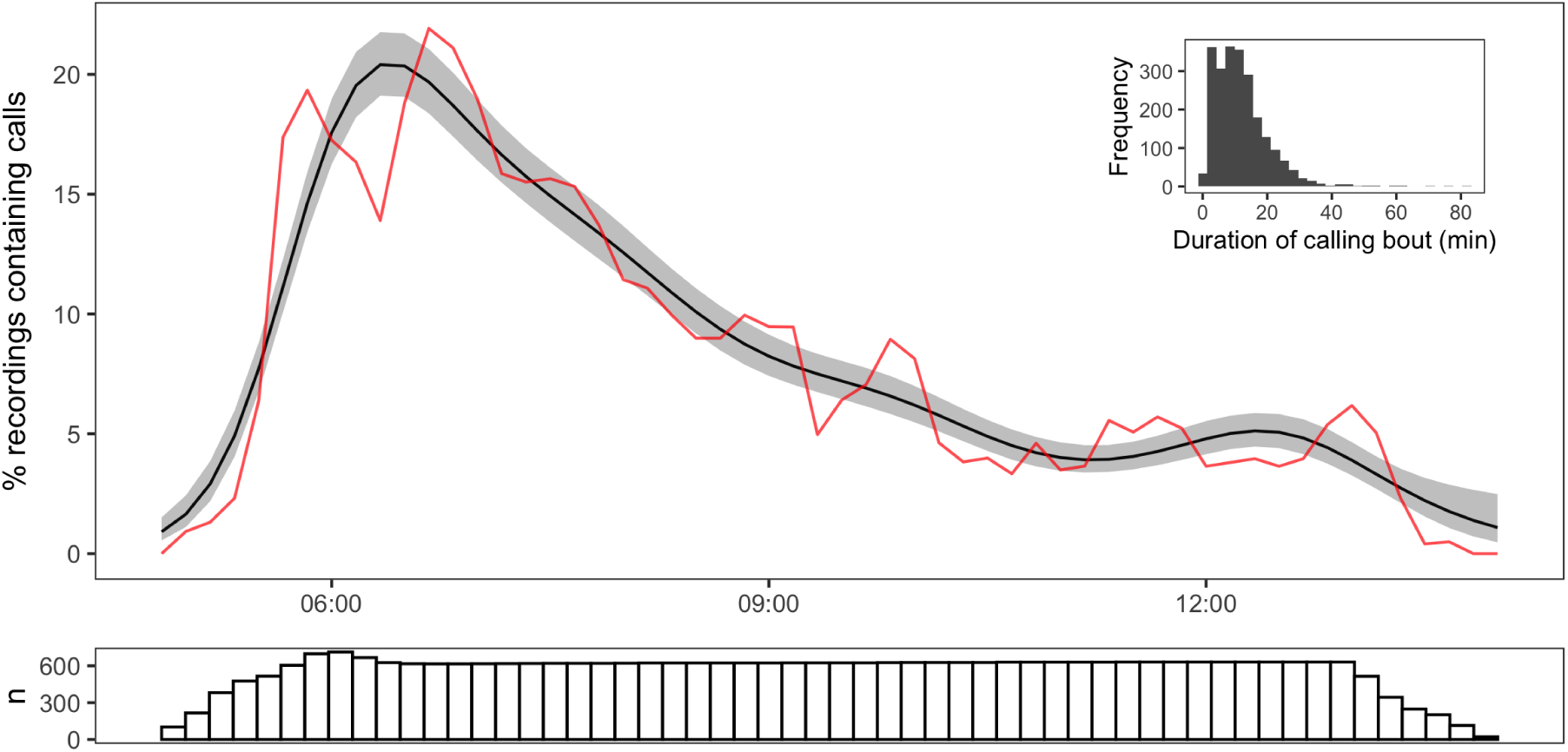
Daily patterns in gibbon calling activity. The red line denotes, per 10 minutes, the proportion of recordings across all locations in which a call was detected (e.g. 05:00-05:10, 05:10-05:20, …). The black line smooths the observed proportions using a GAM (see Supplementary Material D for details). The bottom plot shows the number of recordings per 10-minute segment, showing the survey effort from 05:00–14:00. Peak activity occurs shortly after dawn, dropping rapidly but with some calling activity recorded throughout the morning. Plot inset shows the duration of independent call bouts detected by the classifier. Call bouts are intervals of regular calling, with no detected call 200s either side of the bout. Daily calling typically consists of a number of calling bouts.

## 5 Discussion

Long-term monitoring will generate thousands of hours of recordings across multiple survey sites, and manually labelling these recordings is typically infeasible given logistical constraints. Our results demonstrate that passive acoustic monitoring incorporating an automated classifier can be an effective tool for remote detection of calling species, potentially enabling systematic monitoring whilst saving time, funds and manpower. Our approach, applied to Hainan gibbons, is general and easily extended to other calling species.

Our models allow new recordings to be classified on a per-second basis, to a high degree of accuracy. Although perhaps false negative rates of 1.7% may not be sufficiently low for full automation of Hainan gibbon call monitoring, they greatly facilitate the process of manually annotating these datasets by ruling out large portions of recordings that have a near-zero probability of containing gibbon song. In our test datasets, this reduced the amount of audio to be manually processed by 95%. Our model clearly detected all calling bouts in the test data, at the cost of two false positives.

Where false negatives are particularly costly, this is easily incorporated by lowering the threshold required for manual verification. We expect that with more, and more diverse, training data, error rates would decline further.

Where environmental conditions were similar to those used to train the model, predictions were almost perfect and could be used to identify start and end times of call phrases and bouts, returning almost identical values to a human observer. It is impossible to know in advance whether environmental conditions are similar enough to warrant confidence in the associated predictions, but these results suggest that, as more training data covering a range of environmental conditions are added, model applications may go beyond gibbon detection, by automatically extracting inputs for more detailed behavioural analyses, for example of gibbon call syntax (Clarke, Reichard, & Zuberbühler, 2006).

Practically, developing an acoustic classifier such as ours requires a number of steps: deciding on an appropriate unit of analysis; manually labelling data; augmenting data and allocating it between training, validation, and test sets; choosing and fitting appropriate neural network models; and selecting a preferred model and using it to process the unlabelled portion of the data. Our study illustrates how model development and implementation are informed and guided by ecological objectives, here primarily detecting gibbon vocalizations over time scales of minutes or hours, and domain knowledge of Hainan gibbon call behaviour.

We based our classifier on phrases, rather than shorter notes or longer calling bouts, to balance ease of identification with data availability and computational requirements. Individual notes are easily confused with other sources (see Figure 2b). While calling bouts are highly distinctive, there are relatively few of them and, being longer in duration, they require more parameters to capture the same degree of detail. Phrases are far more numerous, less variable, and require fewer parameters.

Given this choice, segment duration was chosen to be longer than the longest phrase across all training data (8 seconds). The slightly longer segment length provides more presence segments – for example, an 8s phrase results in three 10s presence segments, but would only result in a single segment if the segment length was restricted to 8s. Preliminary runs based on shorter segments of 0.5–2 seconds and *partial* phrases did not yield good performance, with many false positives, probably because a small segment is not enough to distinguish gibbons from other species calling within the same frequency range.

Even using phrases, we have relatively few positive examples and these occur within a highly variable background environment, which is likely to be a common situation for ecological applications. The amount of data available to train neural networks is important, and CNNs tend to require relatively large amounts of data (at least thousands of each class) to generalize well. It may often be possible, as in our case, to collect or label additional data, but data augmentation is a valuable low-cost strategy for increasing sample sizes in conjunction with these other more effort-intensive approaches (Bergler et al., 2019; Hestness et al., 2017; Kahl et al., 2017; Sun et al., 2017). In practice the process can be an iterative one guided by subjective judgement. We initially annotated only 40h across five recordings, but models based on these were poor, even with augmentation. Model performance (on the same test set) improved as we add more training data; we were also able to create more complex neural networks. Gains in accuracy decreased with additional annotations, and we stopped when these became marginal, but presumably further increases are possible as novel environments are included.

Training, validation and test datasets should be constructed by allocating longer contiguous sequences of audio to each of these, and then preprocessing each of these, rather than randomly allocating the segments themselves, which are highly autocorrelated and will thus overstate test accuracy. Wherever possible, we recommend using entirely independent recordings in the test dataset.

We found that 2-D CNNs based on spectrograms performed substantially better than 1-D CNNs that use amplitude time series following some initial preprocessing, mirroring Stowell, Wood, et al. (2019). Deep neural networks are often motivated by an argument that they learn salient features, rather than having to have these provided to them, but where intermediate features (here, spectral densities) can be provided, these speed up the learning process and provide measurable benefits. Beyond the 2-D/1-D distinction, we found that network architectures had relatively little impact on model accuracy, and we achieved good performance using relatively small, simple network architectures, again motivated by limitations on training data. We used few dense layers, each with only a small number of nodes, as these are particularly parameter hungry. Our basic approach was to start with simple architectures, evaluate them, and then add complexity in an iterative manner. Traditional performance metrics such as precision and recall, while important, are not the only relevant measures of classifier success. Practically, classifiers such as ours can be used to point to audio segments that possibly contain gibbon calls, and that require manual verification. Where classification accuracy lags behind that of human experts, or where errors are costly – that is, in many ecological applications – attention shifts from replacing manual annotation to facilitating it. Probability cutoffs can be calibrated to balance the costs of false positives and negatives, and, even if the model is wrong by a few seconds, the amount of time spent in manual verification, compared to that required to processing the entire file manually, is minimal. Our classifier reduces an eight-hour recording to on average 22 minutes with false positive and negative rates under 2%. This time can be further reduced by playing back only those 10s segments that are predicted to contain phrases, although in our case the reduction in overall time was offset by the difficulty of manually verifying segments that are often not contiguous in time.

Analysis of our multi-month dataset demonstrated that gibbons could be detected regularly across all selected survey points, with call detection consistent with known patterns of gibbon behaviour and ecology. Calls were detected at expected times (Chan et al., 2005), and our dataset provides a more precise baseline on Hainan gibbon call timing and duration. Hainan gibbon calling bouts were also generally detected less frequently during the wet season, a period when other gibbon species are also known to sing less frequently (Cheyne, 2008; Clink, Ahmad, & Klinck, 2020). Interestingly, call bouts recorded within the area occupied by a solitary male gibbon were amongst the shortest recorded bouts, and started and finished later than bouts from known social groups. While we cannot exclude the possibility of detecting group calls at this location, this finding suggests important new information on the behavioural ecology of solitary Hainan gibbons that may assist future monitoring and conservation planning.

It is uncertain whether within-recorder and between-recorder variation in calling bout detections represents variation in calling frequency between groups, and/or variation in detection effectiveness by recorders, with the latter possibility likely associated with specific recorder placement, local terrain, specific gibbon movement patterns across landscapes, and group home range size (cf. Bryant et al. (2017)). Future work could investigate detection likelihood in relation to specific environmental parameters and local weather conditions (e.g., rainfall, wind, temperature), data on which were not available for our survey period but are known to affect calling behaviour in other gibbons (Coudrat, Nanthavong, Ngoprasert, Suwanwaree, & Savini, 2015; Yin et al., 2016).

Where calls can be detected across multiple recording locations, acoustic spatial capturerecapture methods provide a means of estimating animal abundance (Stevenson et al., 2015). While our locations are too far apart for this to be feasible, this represents an important next step in monitoring a critically endangered population. Classifiers capable of discriminating between groups or individuals can be valuable inputs to this process (Augustine, Royle, Linden, & Fuller, 2020), as well as providing insight into the behavioural ecology of groups or individuals. We also recommend that call detection ranges should be determined for the specific field conditions at BNNR (e.g., slope, vegetation density), to calibrate monitoring effectiveness of specific recorders, and determine effective recorder placement (grid area/density) to ensure saturation of monitoring coverage. However, passive acoustic monitoring can now be introduced as an important component of the Hainan gibbon conservation toolkit, both for future use at BNNR and also to potentially detect unknown remnant gibbon populations elsewhere across Hainan (S. T. Turvey et al., 2017). Our classifier permits rapid and potentially real-time monitoring of Hainan gibbons, and we hope that the approach we describe in developing this classifier can serve as a roadmap for practitioners to implement their own classifier for other passive acoustic monitoring projects, and contribute to the effective conservation of calling species.

## Acknowledgements

We thank the Management Office of Bawangling National Nature Reserve for logistical assistance in the field. Fieldwork was funded by an Arcus Foundation grant to STT and a Wildlife Acoustics grant to JVB. ID is supported in part by funding from the National Research Foundation of South Africa (Grant ID 90782, 105782). ED is supported by a postdoctoral fellowship from the African Institute for Mathematical Sciences South Africa, Stellenbosch University and the Next Einstein Initiative. This work was carried out with the aid of a grant from the International Development Research Centre, Ottawa, Canada (www.idrc.ca), and with financial support from the Government of Canada, provided through Global Affairs Canada (GAC; www.international.gc.ca). We also thank the following rangers who contributed to data collection: Guang Wei, Zhong Zhao, Qing Lin, Jinbing Zhang, Zhicheng Zhang, Quanjin Li, Xiaoliang Fu, Zhengchong Zhou, Lubiao Huang, Zhengkun Ye, Zhenghai Zou, Jinqiang Wang, Wentao Han and Zengnan Xie.

## Authors’ contributions

ST, JB and HM conceived the passive monitoring project and developed the study designs and protocols. ED, ID and JH conceived the development of an automated classifier and designed the methodology. WL, ZL, QC, ZZ, HM and JB were responsible for fieldwork and data collection. ED, AH and JB annotated the data. ED constructed the classifier and performed the analysis. ED, ID and ST wrote the paper. All authors contributed critically to the drafts and gave final approval for publication.

## Data accessibility

All code for training and testing the neural networks and conducting additional analyses is available at https://github.com/emmanueldufourq/GibbonClassifier (Supplementary Material E). A subset of acoustic recordings, including training and testing labels, has been stored on Zenodo: https://doi.org/10.5281/zenodo.3991714.

## Supplementary Material

### A Details of observed call bouts in training data

In the preliminary stage of model building we used a subset of 72 hours of recordings (nine eight-hour recordings) to inform our decision to use a window of 10s. Across these recordings, an average of 2.3 calling bouts were observed per eight-hour period (min 1, max 4), with on average 54 phrases per bout (min 31, max 116). The average duration between phrases within a calling bout was 19.4s. Table A.1 presents the distribution of the numbers of syllables per phrase, as well as the mean duration of phrases consisting of different numbers of syllables. All phrases contained between one and six phrases, with the majority of phrases made up of one to four syllables.

**Table A.1:** The average total duration for each type of hainan gibbon song. These are the syllables in the long calls that the hainan gibbon’s perform. The number of times each type occurs is also presented. These values also include the breaks between consecutive calls.

| Type | Average total duration | Number of instances |
| --- | --- | --- |
| 1 syllable | $2.6 \pm 0.9$ | 175 |
| 2 syllables | $4.3 \pm 0.8$ | 413 |
| 3 syllables | $5.1 \pm 0.9$ | 339 |
| 4 syllables | $5.8 \pm 0.8$ | 302 |
| 5 syllables | $6.4 \pm 0.8$ | 62 |
| 6 syllables | $6.0 \pm 0.6$ | 13 |
| Duet | $6.1 \pm 0.9$ | 56 |

### B Details of predicted call bouts in test data

Table B.1 shows observed and predicted start and end times of calling bouts in nine eight-hour recordings used to test our final (2-D CNN) model. Each bout is denoted by [*t*_*s*_, *t*_*e*_], where *t*_*s*_ and *t*_*e*_ are start and end times (in seconds from the start of the recording) respectively. No calling bouts were missed, but two predicted bouts were false positives (denoted in bold) - these are 52 and 272 seconds of false positives respectively.

### C Seasonal differences in gibbon detections

Table C.1 reports the same summary statistics as Table 2 in the main text, but separately for wet and dry seasons. Gibbons called substantially less frequently in the wet season at four sites (2, 5, 8), more frequently at two sites (4, 7), and less on average across all sites (65% (261/403) vs. 77% (305/398)). Calling occurred over a subtantially greater part of the day in the wet season (07:05–10:25) than in the dry season (07:25–09:30), although mean calling time per day did not differ substantially (wet season = 30m, dry season = 29m).

**Table B.1:** Observed and predicted start and end times (sec) of calling bouts

| File | Type | Bouts (seconds) |
| --- | --- | --- |
| 1 | Correct | [3682 3899], [3911 4174] |
|  | Predicted | [3357, 4180] |
| 2 | Correct | [3349 3831], [8854 9456], [14796 15502] |
|  | Predicted | [3342, 3836], [8850, 9459], [14791, 15506] |
| 3 | Correct | [3676 3795], [14759 14955], [19557 20257], [20533 20856] |
|  | Predicted | [3623, 3802], [14752, 14962], [19365, 20262], [20526, 20860] |
| 4 | Correct | [3950 4201], [5390 5941] |
|  | Predicted | [3945, 4208], [5351, 5948] |
| 5 | Correct | [3398 4148], [8507 9018], [10642 11035], [14918 15542] |
|  | Predicted | [3366, 4154], [8477, 9024], [10509, 11039], [14911, 15548] |
| 6 | Correct | [3423 3783], [6370 7086] |
|  | Predicted | [1216, 1268], [3417, 3789], [6367, 7091] |
| 7 | Correct | [5607 6626] |
|  | Predicted | [1704, 1976], [5627, 6629] |
| 8 | Correct | [3133 3802], [11643 12317] |
|  | Predicted | [3312, 4028], [11488, 12322] |
| 9 | Correct | [10210 10235], [24377 25125] |
|  | Predicted | [10184, 10239], [24373, 25129] |

**Table C.1:** Detection of gibbon calling bouts by different recorders in wet and dry seasons.

| Location | Survey days | % days calls detected | Mean calling time per day (min) | Mean start time of first bout | Mean end time of last bout |
| --- | --- | --- | --- | --- | --- |
| **Dry season (Mar-Apr), 61 days** |  |  |  |  |  |
| 1 | 52 | 75 | 28.7 | 07:19 | 09:26 |
| 2 | 37 | 73 | 26.4 | 07:00 | 08:48 |
| 3 | 58 | 81 | 34.8 | 08:01 | 10:09 |
| 4 | 52 | 81 | 28.6 | 07:46 | 09:25 |
| 5 | 51 | 51 | 29.9 | 07:31 | 09:23 |
| 6 | 55 | 80 | 28.3 | 08:04 | 10:10 |
| 7 | 46 | 78 | 21.6 | 07:07 | 08:46 |
| 8 | 47 | 94 | 31.2 | 06:33 | 09:26 |
| **Wet season (May-Aug), 100 days** |  |  |  |  |  |
| 1 | 35 | 63 | 16.2 | 08:00 | 10:09 |
| 2 | 53 | 26 | 36.8 | 06:54 | 09:58 |
| 3 | 45 | 82 | 26.7 | 06:49 | 10:24 |
| 4 | 53 | 91 | 24.5 | 07:42 | 10:16 |
| 5 | 28 | 0 | — | — | — |
| 6 | 48 | 77 | 19.9 | 07:48 | 10:20 |
| 7 | 83 | 95 | 35.2 | 06:47 | 10:25 |
| 8 | 58 | 41 | 58.5 | 05:45 | 11:05 |

### D Generalized additive model details

We fitted a generalized additive model (GAM) with the *mgcv* package in R (Wood, 2017) to model the relationship between the number of detected gibbon call bouts and time-of-day. A binomial distribution for the error terms and an log link function was used, with a smooth term using cubic regression splines with 10 knots (*k* = 10) capturing non-linearities in the relationship between predictor and response variable. The exact number of knots is not critical but was chosen conservatively with the intention of producing biologically meaningful results. We checked that we did not over-specify the number of knots using the effective degrees of freedom as a guide. The model explained 88% of the variability in detected counts (deviance explained) and residual analysis plots indicated symmetrically distributed residuals. There was no discernible evidence of heteroskedasticity or unmodelled relationships between residuals and either observed or fitted values of the dependent variable.

### E Software

Two interactive notebooks, *Train.ipynb* and *Predict.ipynb*, illustrate the two main processes in developing an automated classifier: pre-processing audio file and training a convolutional neural network (*Train.ipynb*) and using an already-constructed model to identify calls in a new and unlabelled recordings (*Predict.ipynb*).

A detailed manual is provided in the same repository as the code, so here we only briefly illustrate the workflow (Figure E.1). Users first need to download the code repository and install all requirements in *requirements.txt* using pip install -r requirements.txt.

For training, input data takes the form of (a) one or more .wav files containing already annotated recordings, (b) a text file containing the annotated call times in the training files, and (c) a text file containing the filenames of these .wav files. Upon downloading the repository, an example of (a) and (c) is downloaded to the *Raw_Data/Train* and *Call_Labels* folders, while an example of (b) appears as *Training_Files.txt* in the root directory. Folders and filenames can be changed as notebook options, as well as optional parameters controlling various aspects of model building (downsampling rate, augmentation, etc). Two core functions execute_preprocessing_all_files and train_model perform preprocessing (creating image files containing spectrograms) and build the CNNs.

For predicting on test or unlabelled files, users specify the location of the test .wav file, as well as the location of the model parameters obtained during training. Model weights for our best 2-D CNN are downloaded with the repository and saved as *Experiments/pretrained_weights_from_paper.hdf5*, so this notebook will run directly on new data without needing to retrain the model. The function execute_processing runs the test file through the trained neural network and outputs predicted call times as a spreadsheet.

**Figure E.1:**
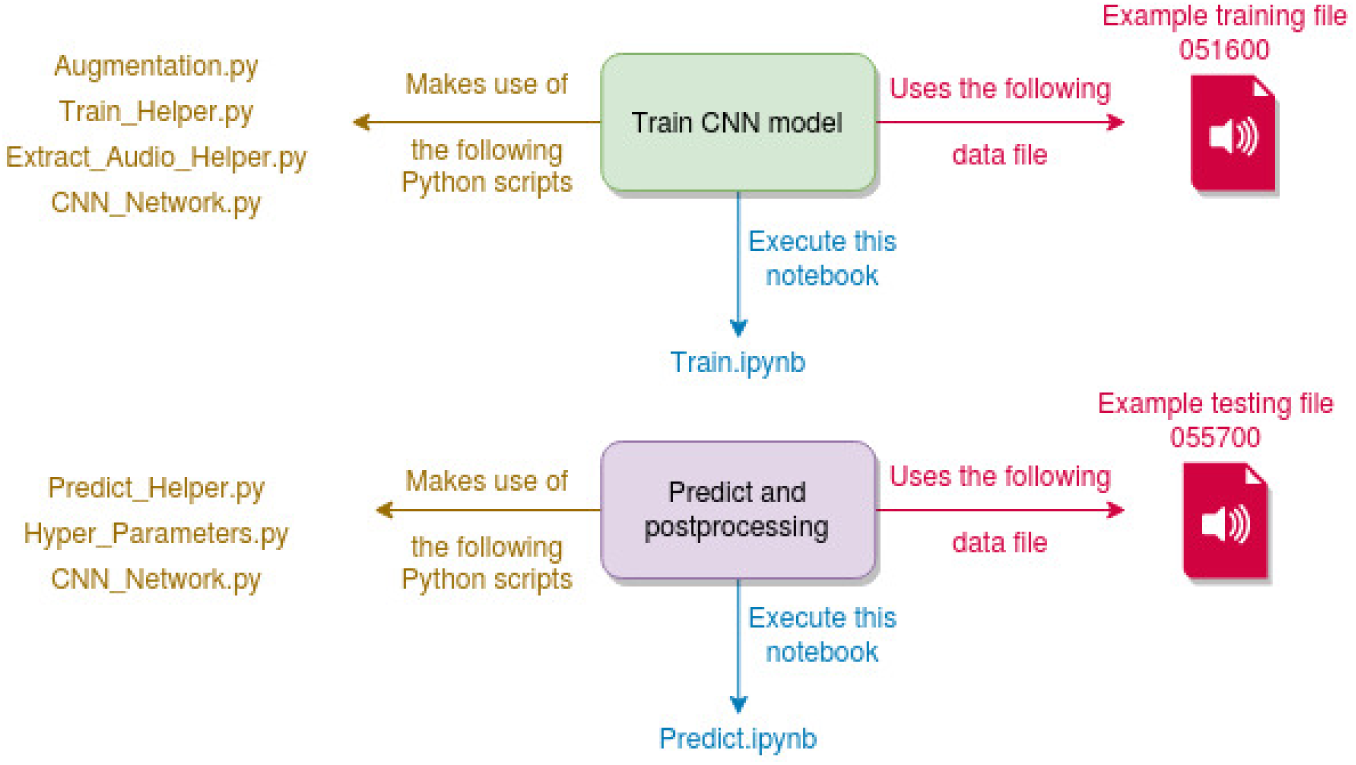
Illustrating the pipeline and code dependencies for training and prediction.

